# Skew in ovarian activation depends on domicile size in a facultatively social thrips

**DOI:** 10.1101/038414

**Authors:** J. D. J. Gilbert, A. Wells, S. J. Simpson

## Abstract

Costs and benefits of group living are a fundamental topic in behavioural ecology. Resource availability affects individuals breeding prospects alone and in groups, as well as how reproduction is distributed within groups (“reproductive skew”). Here, we provide correlational evidence in facultatively social thrips that breeding resources are associated with (1) whether solitary or social living is favoured, and (2) the degree of ovarian skew.

*Dunatothrips aneurae* (Thysanoptera, Phlaeothripidae) cooperatively build silk “domiciles” on Australian Acacias, feeding exclusively from internal phyllode surfaces. *Per capita* productivity scaled differently with group size depending on domicile volume – females in small domiciles did better alone than in groups, whereas in large domiciles single and group-nesting females did equally well. Ovarian dissections revealed that in small domiciles some females were nonreproductive, indicating ovarian (i.e. reproductive) skew. Skew increased as domicile size decreased and group size increased. Breeders had smaller oocyte volume in smaller domiciles, especially those containing nonreproductives.

These findings suggest group formation and reproductive skew in *D. aneurae* may be influenced by reproductive competition for breeding resources. Nonreproductive females in small domiciles may be reproductively suppressed, subfertile, or waiting to reproduce. We speculate they may avoid eviction by contributing as “helpers” to domicile maintenance.

## INTRODUCTION

Should animals breed alone or in groups? The size of animal breeding groups represents the balance of multiple benefits and costs to individuals [reviewed in 1]. From the perspective of costs, groups may form because costs of independent breeding outweigh costs of grouping [2]; such “ecological constraints” include, for example, habitat saturation in vertebrates, or nest construction in invertebrates [3-7]. Viewed the other way, the benefits to the individual of being in a group may outweigh the benefits of independent breeding [8]; for example, groups maybe better than individuals at securing high-quality resources [e.g. 9]. In general, breeding success depends upon access to a sufficient amount and quality of resources – whether individually [e.g. 10] or within groups [e.g. 11,12]. We would therefore expect the size and quality of resources available to individuals inside and outside of groups to affect the relative costs and benefits of group breeding [13,14].

Within a group, though, an individuals breeding prospects are often far from guaranteed. Reproduction can be shared unequally – to an extent that is termed the reproductive skew [15]. While low-skew societies are relatively egalitarian, in societies with high skew, fully reproductive breeders exist alongside subordinate individuals that breed to a lesser extent or not at all, who may or may not assist breeders [e.g. 16,17,18]. Competition for resources within groups may affect reproductive skew [e.g. 12,reviewed in 19]. Thus, we might expect resource size and quality to be associated not only with group size, but also with reproductive skew.

We might intuitively expect groups inhabiting bigger or better resource patches to have lower skew; for any given group size, a better resource patch can support a higher proportion of individuals reproducing. Additionally, better resource patches can also support larger groups – which may have lower skew irrespective of resources, for three reasons. First, larger groups may be harder for a dominant individual to control [19]. Second, in systems where group reproduction is directly related to subordinates effort, subordinates in larger groups may contribute relatively less to group productivity compared to their expected success when breeding alone – and so may demand a greater share of reproduction as a “staying incentive” [20]. Finally, in larger groups there may be a greater threat of aggressive challenges for dominance, such that dominant individuals may be more motivated to concede reproduction to subordinates as a “peace incentive” [21,22].

Acacia thrips (Thysanoptera, Phlaeothripinae) include species with a range of social behaviour. As haplodiploids, they present contrasts and comparisons with the better-studied social Hymenoptera, and are increasingly appreciated as a parallel model clade for social research [23]. Two lineages of Australian Acacia thrips exhibit social behaviour. In both lineages, group members live, feed and reproduce entirely within their resource. In gall-inducing thrips, which are eusocial, a general evolutionary trend towards smaller galls (i.e. resources) has been accompanied by more pronounced division of reproduction between foundress and soldiers, i.e. increasing reproductive skew [24-26]. This finding was interpreted as evidence that within-gall competition for resources may have helped to drive the evolution of the soldier caste [24]. Yet, without a truly solitary option available, the effects of resource size upon the costs and benefits of social behaviour are hard to assess.

The other lineage contains several origins of “phyllode-glueing”, a much less well-studied lifestyle. Phyllode-gluers live and breed entirely within “domiciles” constructed by glueing together Acacia phyllodes with a silk-like cement extruded from their abdomen [23]. Among these, a few species in the genus *Dunatothrips* show facultative pleometrosis, i.e. joint nesting. Females of some species are solitary (e.g. *D. gloius)*, but in others can be found in groups of up to 4 in *D. skene* [27], 8 in *D. armatus* [23], ∼15-20 in *D. aneurae* [27-29] and >70 in *D. vestitor* [23].

By far the best studied of these species is *D. aneurae*. In this species, single foundresses comprise roughly 70% of the population [27,28,30], showing that independent nesting is certainly feasible, and suggesting that ecological constraints driving social behaviour are weak. Furthermore, foundress numbers were found not to be correlated with domicile density on a tree, suggesting that habitats are not locally saturated [28]. Nevertheless, per capita productivity appears to decline with increasing foundress numbers [28], suggesting a cost of grouping. Several possible benefits of joint nesting may counterbalance this cost. Cofoundresses tend to be relatives, enhancing inclusive fitness of group members [30]. Cofounding enhances defence against kleptoparasites [28] although not against inquilines [31], and also increases adult survival via other unknown means [28] – hypothesized to involve sharing of costs associated with e.g. domicile building or maintenance [23,29,32].

Variation in resources available to single versus multiple females may affect any of these costs and benefits – and ultimately whether social behaviour is favoured. Indeed, Crespi et al. [23] made the suggestion that resource variability may be one key reason why *D. aneurae* and *D. vestitor* may have evolved to have such a high degree of social flexibility. Domicile sizes in these species are highly variable because of the loose, multi-phyllode conformation of their domiciles compared to more solitary species, which tend simply to make a domicile in the diamond-shaped space created by two crossed phyllodes. In this study, we aim to elucidate whether the size of *D. aneurae* domiciles, i.e. of breeding resources (feeding area or breeding space), is associated with per capita reproduction and its distribution within a domicile (reproductive skew).

## METHODS

*Dunatothrips aneurae* domiciles occur predominantly on terminal phyllodes of narrow-phyllode varieties of *A aneura* [29]. Their principal function is to reduce desiccation in the arid environment [32]. Domicile construction is aseasonal, and appears to require male presence [29], although founding males are seldom found in field domiciles (Bono & Crespi, 2006). After construction, *D. aneurae* foundresses cast off their wings and produce one generation of offspring (Morris et al., 2002). Most offspring disperse but indirect evidence suggests some may reproduce in the natal domicile (Bono & Crespi, 2008). Domiciles may be extended over time [33] although most are simple and consist of a single chamber (Gilbert & Simpson, unpublished data).

In fieldwork trips between September 2011 and October 2013 we collected 513 *D. aneurae* domiciles from thin-leaved variants of *A aneura* on the Fowlers Gap property, approx. 110km N of Broken Hill, Australia [see 29 for location details] and placed them in a fridge at 4°C until dissection under a binocular microscope. Most domiciles were dissected within 3 days of collection; 91 domiciles were placed in a freezer at −20°C and dissected approx. 1 year later (excluding these domiciles from analysis had no appreciable effect upon results).

### Domicile volume, foundress number and per capita productivity

We measured domicile volume as a simple cuboid, i.e. length × width × depth at their largest dimensions, ignoring any convolutions. Most domiciles were of a reasonably simple shape that warranted this assumption.

We counted domicile inhabitants and classified them as foundresses [dealate females; 28,29], offspring of all stages up to alate adults, or, rarely, adult males. Adult males do not lose their wings upon reproduction and so may potentially be confused with adult offspring. However, adult males are very rare in active *D. aneurae* domiciles [27-29]. Accordingly, any single adult males that occurred alongside alate female adults, plus multiple adult males occurring in the same domicile, were assumed to be male alate offspring. Single adult males found in domiciles with only dealate females and eggs or very young larvae present were assumed to be reproductive males and excluded from analysis. We also counted eggs and classified them as hatched or unhatched. We excluded from analysis any domiciles that were under construction, which contained no offspring and/or no foundress, and domiciles with only adult offspring (from which some offspring may have already dispersed).

We calculated total domicile offspring as the sum of the number of live offspring in each domicile and the number of unhatched eggs present. We did not attempt to count failed offspring within nests, nor the probability of whole-nest failure. Per capita offspring was calculated simply as the total offspring divided by the number of foundresses. We analysed per capita offspring as a function of domicile volume and foundress number using a generalized linear model (GLM) with a Poisson distribution.

### Ovarian status

In a subset of domiciles (n_domiciles_=164, n_foundresses_ =267), foundresses were then killed by immersing in 100% ethanol for 1 minute, and were dissected immediately in water. We measured the pronotum width using an eyepiece reticle. The extent of ovary development could be clearly classified as developed or undeveloped (Figure 1). In developed ovaries, we measured the length and width of any developing oocytes. The volume of each oocyte was calculated as an ellipsoid (π × length × (width/2)^2^) and the resulting volumes were summed to give the total volume of developing oocytes in each ovary.

**Figure 1.**
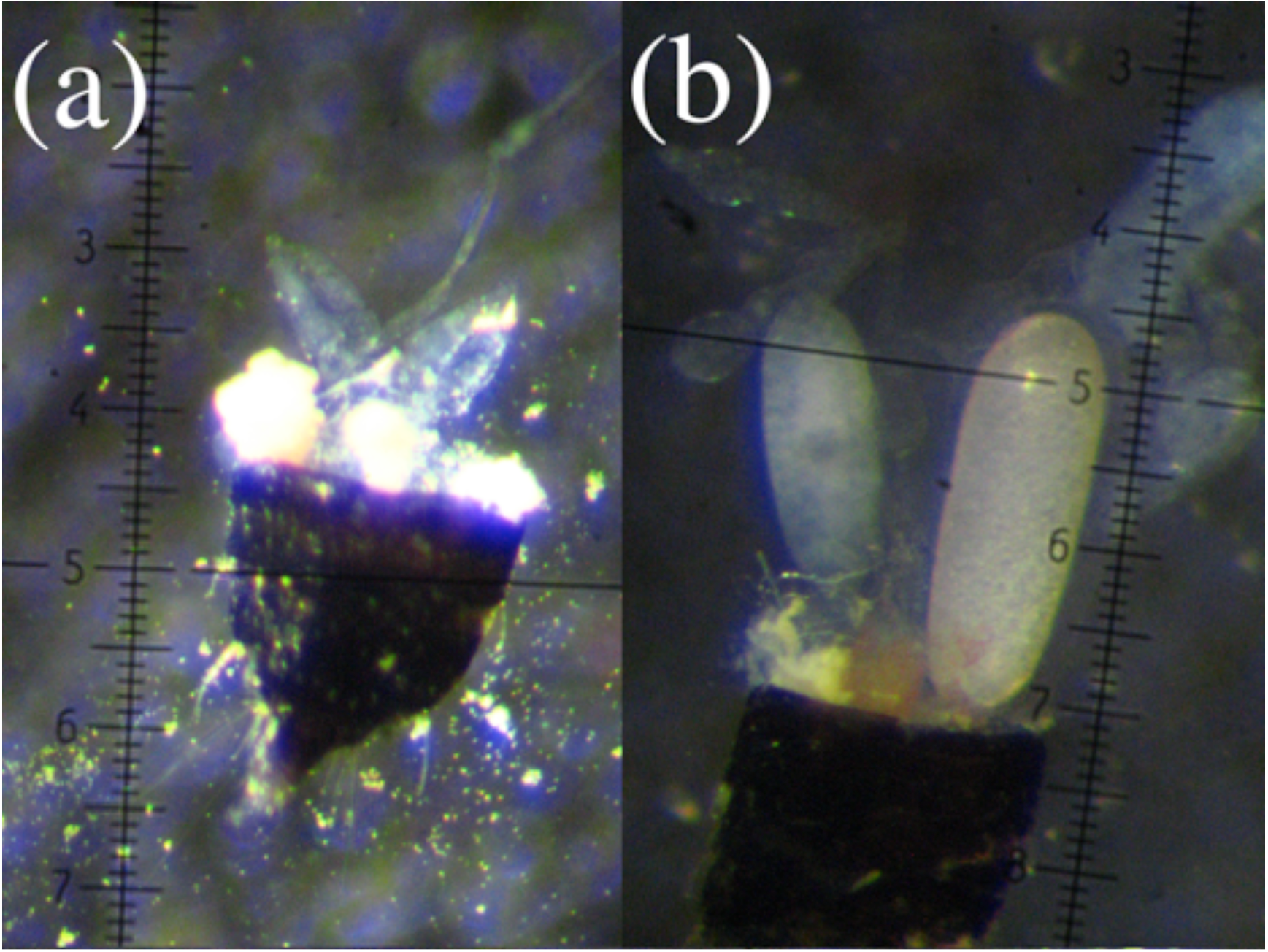
Dissected ovaries of *Dunatothrips aneurae* foundress females: (a) ovary with no developing oocytes (50x; 1 reticle unit = 20 um); (b) ovary with mature oocytes (50x).

We analysed ovarian status (developed or undeveloped) as a function of domicile volume, foundress number and body size (pronotum width) using a generalized linear mixed model (GLMM) with a binomial distribution and “domicile ID” as a random factor. The probability of a given female being nonreproductive (i.e. with no developing oocytes) can be viewed as an estimate of skew – although note that estimates based on ovarian dissections represent a conservative minimum estimate. Genetic data would be necessary to test whether those females that do have developing oocytes are actually ovipositing within the nest [see e.g. 26,34,35].

Among the subset of females that were reproductive, we analysed the total volume of developing oocytes using a linear mixed model with domicile volume, foundress number, pronotum width and the presence/absence of nonreproductive females as predictor variables and “domicile ID” as a random factor.

## RESULTS

### Productivity

Domicile volume ranged from 7.75 mm3 to 1932.78 mm3 (mean 205.76 ± SE 11.35 [SD 256.29]). Within domiciles, foundress numbers ranged from 1 to 22 (median 2 ± SD 2.72). Per capita productivity (total offspring/number of foundresses) ranged from 0.16 to 19.67 (median 4.33 ± SD 3.80), and was associated with the interaction of foundress number (single vs. multiple) and a quadratic effect of domicile volume (GLM with quasipoisson distribution: χ^2^=12.99, df=1, p=0.029; Table 1). The quadratic volume term indicated that there was an optimal domicile volume for offspring production, but the significant interaction term meant that this optimum volume differed according to whether domiciles were singly or multiply founded. Specifically, singleton domiciles had a smaller optimum volume than multiply founded domiciles. Over a substantial range of domicile sizes, single females had higher expected per capita reproduction than multiple females (Figure 2). In the smallest third of domiciles, per capita offspring decreased when multiple foundresses were present, whereas in medium-sized or large domiciles, per capita offspring did not change with foundress number (Figure 3).

**Table 1.** Minimal model of per capita productivity (total domicile productivity divided by foundress number; generalised linear model with quasipoisson error distribution) according to domicile volume and foundress numbers. For statistics associated with dropping model terms, see text.

|  | Estimate | SE |
| --- | --- | --- |
| (Intercept) | 1.15E+00 | 3.67E-01 |
| Volume | 1.05E-02 | 4.71E-03 |
| Multiple females | 4.91E-02 | 2.20E-01 |
| Volume<sup>2</sup> | -2.51E-05 | 1.20E-05 |
| Volume x Multiple females | -4.43E-03 | 2.43E-03 |
| Volume<sup>2</sup> x Multiple females | 1.19E-05 | 6.00E-06 |

**Figure 2.**
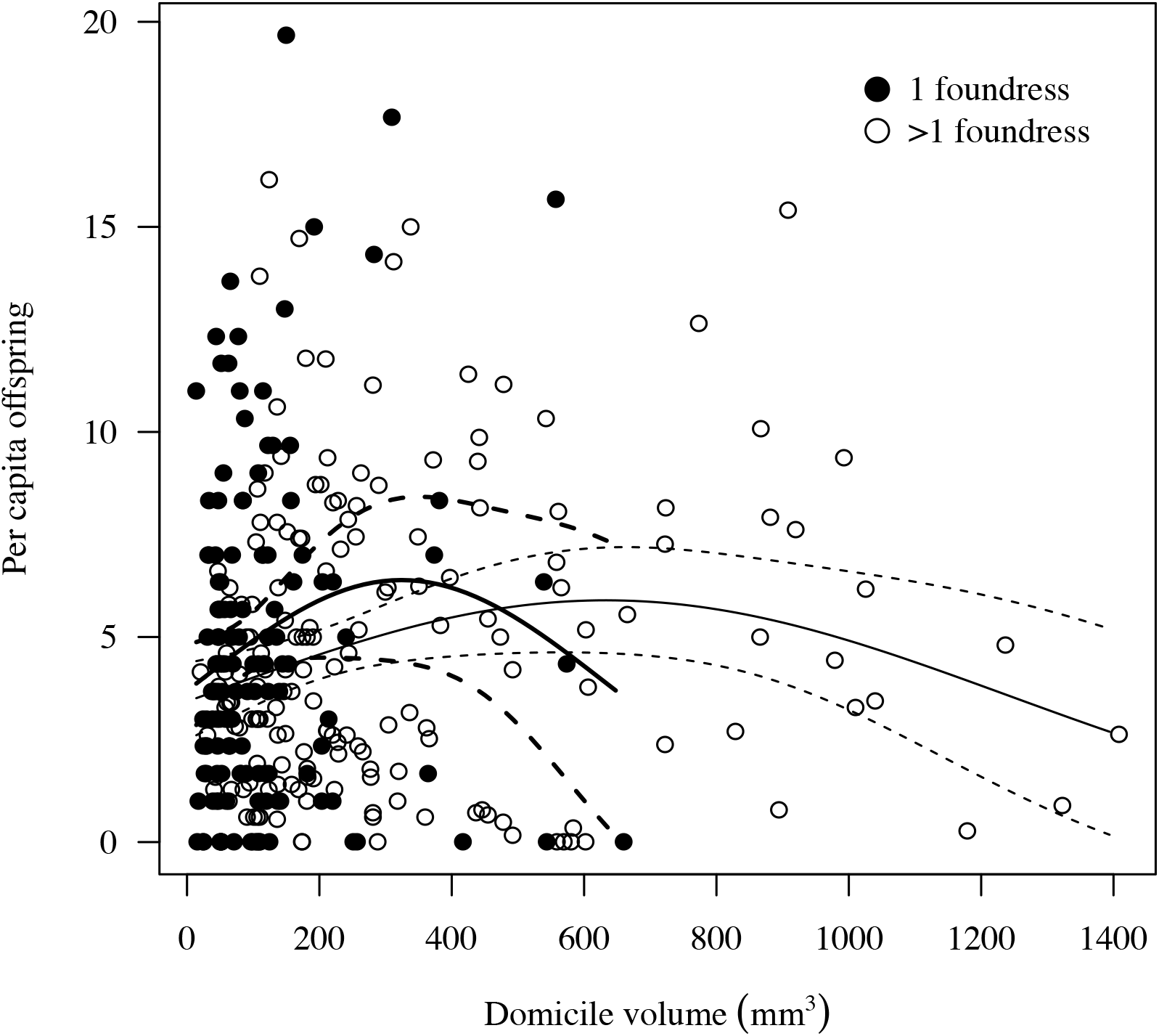
Per capita reproduction according to domicile volume (mm3) in singly and multiply founded domiciles. Best-fit lines and confidence intervals are given from the Poisson GLM.

**Figure 3.**
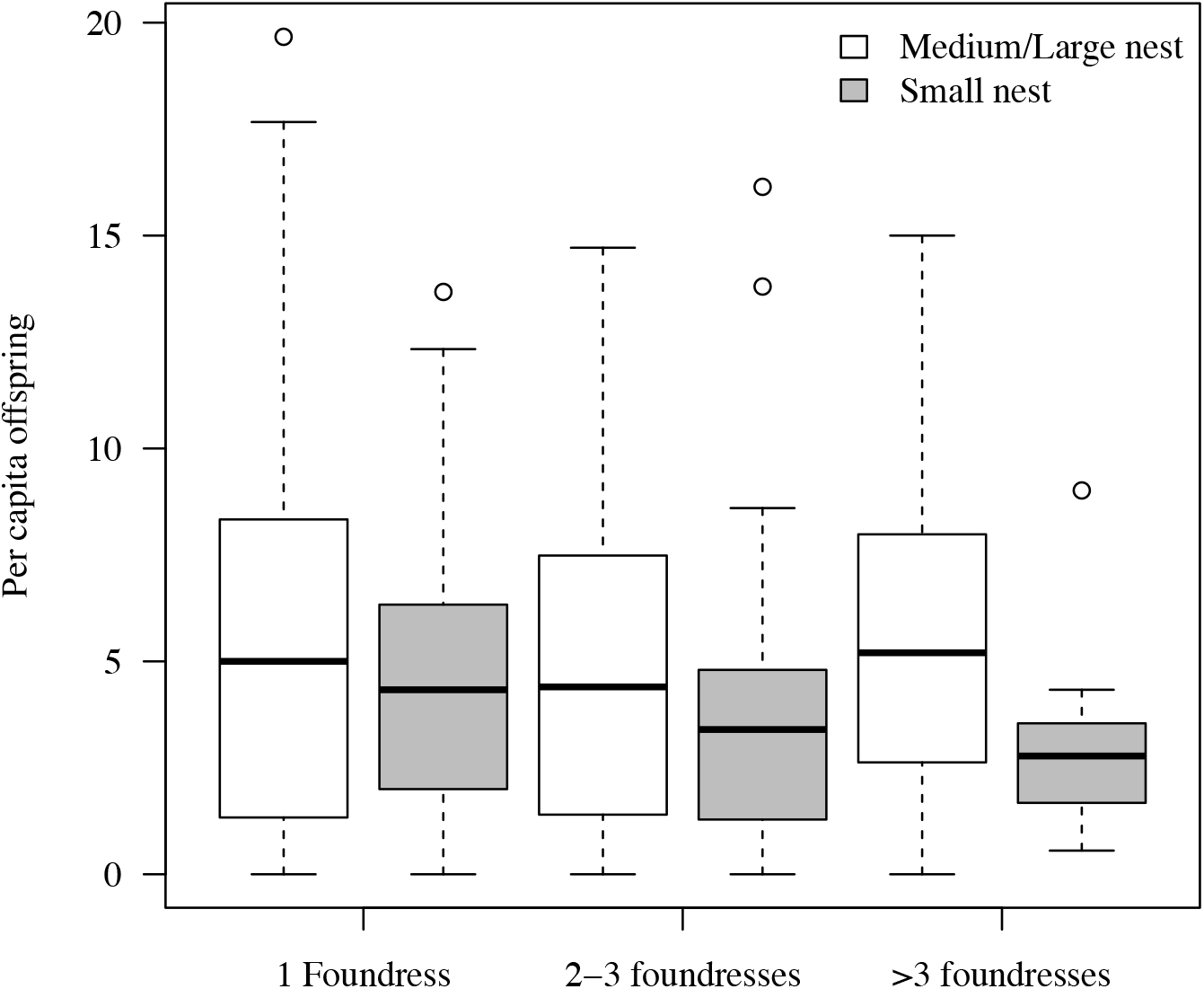
Per capita reproduction in small domiciles versus medium/large domiciles (below and above 33^rd^ percentile of domicile volume, respectively) by different numbers of foundress females.

### Ovarian status

63 out of 267 dissected foundresses (23.5%) had no developing oocytes. Within domiciles, the proportion of nonreproductive females ranged from 0 to 0.8. Among reproductive females, the volume of developing oocytes ranged from 0.39 to 34.9mm^3^.

In the minimal model, foundresses ovarian status was associated independently with both domicile volume (Figure 4a) and foundress number (Figure 4b). Three outliers with very large domiciles were excluded to improve model fit; re-including them gave similar results. Females inhabiting smaller domiciles were generally more likely to be nonreproductive (GLMM with binomial distribution and “domicile id” as a random factor, domicile volume standardized: χ^2^=14.78, df=1, p<0.001; Table 2a). However, females in small multiply founded domiciles were more likely to be nonreproductive than females in small singleton domiciles, whereas in large domiciles almost all individuals were reproductive regardless of foundress number; this was evidenced by a significant main effect of foundress number (χ^2^=5.60, df=1, p=0.018; Figure 3a, b). Larger individuals within associations were not more likely than smaller individuals to have developing oocytes (χ^2^=1.49, df=1, p=0.22). Analysing these data at the domicile level, using a binomial GLM with “proportion reproductive” as the response variable and “domicile volume” and “foundress number” as predictors (but excluding the individual level variable “pronotum width”), resulted in a qualitatively identical minimal model.

**Table 2.** Best models of ovarian status according to foundress numbers and domicile volume: (a) probability that a given female has developing oocytes (generalized linear mixed model with binomial error distribution and “domicile ID” as a random effect; domicile volume standardized); (b) volume of developing oocytes in reproductive females (linear mixed model with “domicile ID” as a random effect). For statistics associated with dropping model terms, see text.

| (a) | Estimate | SE |
| --- | --- | --- |
| Fixed effects |  |  |
| (Intercept) | 3.423 | 0.934 |
| Volume | 1.525 | 0.467 |
| Number of females | -0.525 | 0.240 |
| Random effects |  |  |
| Domicile ID | 2.29 | 1.51 |
| (b) |  |  |
| (Intercept) | 1.321 | 0.193 |
| Volume | 0.0016 | 0.0007 |
| Nonreproductives in domicile | -0.7330 | 0.2874 |
| Random effects |  |  |
| Domicile ID | 0.585 | 0.764 |

**Figure 4.**
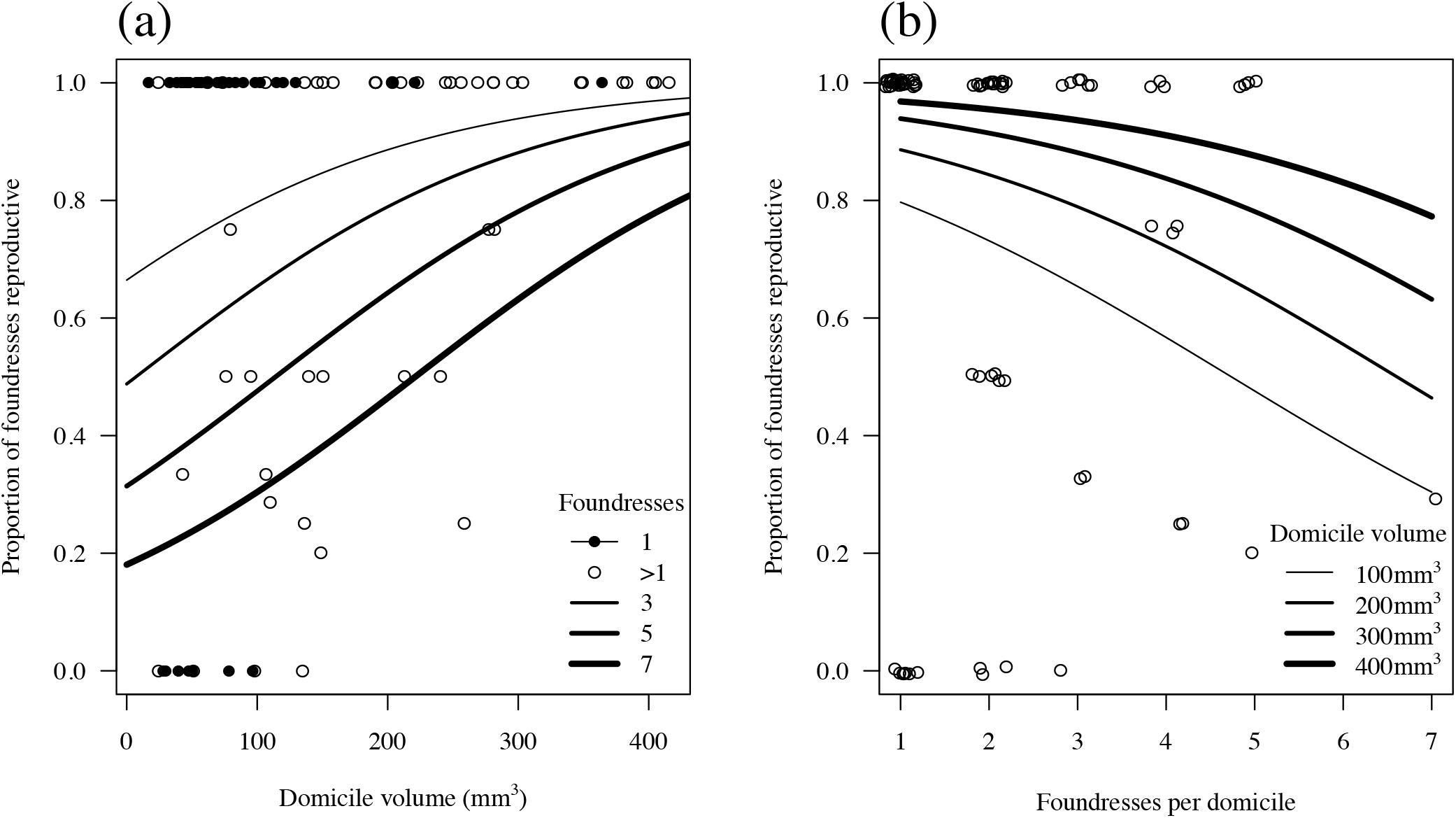
Proportion of reproductive females (i.e. with developing oocytes) according to (a) domicile volume, in domiciles containing different numbers of females; (b) foundress numbers, in domiciles of different volumes. Best-fit lines and confidence intervals are given from the binomial GLM.

Excluding nonreproductives, oocyte volume in reproductive females was smaller in smaller domiciles (linear mixed model with “domicile id” as a random factor, “nest volume” and “pronotum width” scaled, dropping “domicile volume”, χ^2^=4.84, df=1, p=0.027, Figure 5a, Table 2b) and in domiciles containing nonreproductive females (dropping “nonreproductives present in domicile”, χ^2^=6.13, df=1, p=0.013, Figure 5b). Developing oocyte volume did not increase with body size (χ^2^=2.69, df=1, p=0.10) nor with the number of foundresses (χ^2^=0.81, df=1, p=0.37). Again for this analysis we excluded 3 domiciles to improve model fit; re-including them gave similar results. Performing this analysis at the domicile level using a linear model with “mean oocyte volume of reproductive females” as the response variable and “domicile volume”, “foundress number” and “presence of nonreproductive females” (but excluding the individual-level variable “pronotum width”) gave a qualitatively identical minimal model.

**Figure 5.**
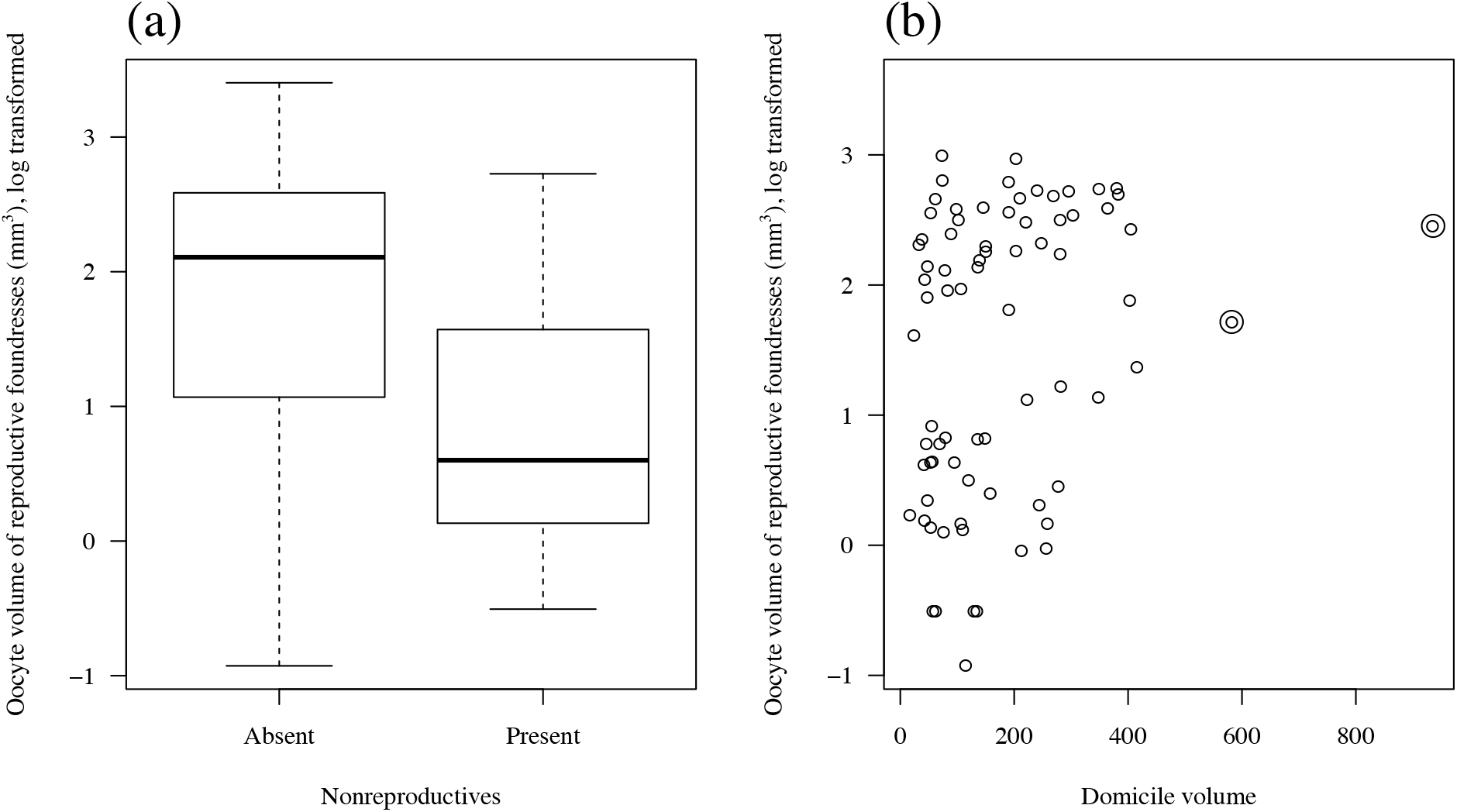
(a) Mean oocyte volume of reproductive females within a domicile (log transformed), with respect to domicile volume. Double open circles are outliers that were removed for the domicile-level analysis; (b) Oocyte volume of reproductive females (log transformed) with respect to the presence or absence of nonreproductives in the domicile.

## DISCUSSION

The relationship between group size and per capita productivity in *D. aneurae* depended upon the size of the domicile: flat in large domiciles, but negative in small domiciles, such that in small domiciles single females had an advantage over multiple females. In smaller domiciles, and in larger groups, an increasing proportion of females were nonreproductive. However, oocyte volume in reproductive females did not decline directly with group size, but instead declined with the presence of nonreproductive females, which in turn tended to occur in smaller domiciles.

Correlational census data must be treated with caution. First, cofounding behaviour may affect the probability of missing data due to whole-nest failure [36,37], something we were unable to determine. Second, the data do not account for the effects of domicile age upon productivity, although we excluded immature and dispersing domiciles as a way of partially accounting for this.

These important caveats notwithstanding, our findings lend support to the idea that resource competition within *D. aneurae* domiciles at least partly determines both per capita reproduction and reproductive skew. This has previously been suggested for Acacia thrips of the eusocial, gall-inducing species [24], which form a sister clade to phyllode-gluers such as *D. aneurae* [23]. In those species, reproductive skew has increased as galls have become progressively smaller over evolutionary time [24,26], suggesting that competition for resources within galls has facilitated the evolution of high reproductive skew (and ultimately of reproductively subordinate soldier castes). By implicitly similar reasoning, Crespi et al [23] suggested that domicile architecture may provide a context for social evolution in *Dunatothrips*. Their basis for this suggestion was that the more-or-less nonsocial species such as *D. gloius* and *D. armatus* tend to construct simple, fairly uniform domiciles out of two crossed phyllodes, while the cofounding species, *D. aneurae* and *D. vestitor*, use many more phyllodes to construct looser, more irregular domiciles with much more size variance; additionally, domicile extensibility may also be important [e.g. 38,39].

In *D. aneurae*, we have shown that similar effects appear to be evident across the range of domicile sizes within a single population. In small domiciles, females did better on their own than in groups – to the extent that some group-living females were actually nonreproductive. By contrast, in large domiciles, individuals did equally well regardless of female numbers, suggesting that competition for resources was not a limiting factor upon reproduction. A similar effect was shown in striped mice [14], which generally live in groups, except in the breeding season, when intense reproductive competition means they form groups only at high population density. Generally, fecundity of plant-exploiting insects can vary with the size or quality of the resource [40]. In social groups, reproduction is often closely linked to resources [12,41] and within-group competition is frequently an agent of reproductive suppression in both vertebrate and invertebrate societies [reviewed in 19,42,43]. In species that cohabit galls, resource competition can limit reproduction for inhabitants [reviewed in 44]. More generally, nest morphology has been implicated in social evolution in a variety of taxa [24,45-48].

Hence it may be that breeding in *D. aneurae* is despotic or communal depending on the extent of competition for resources. The question remains (as for the gall-inducing thrips): by what mechanism does such resource competition operate? For example, the negative association between resource size and skew that we observe here would be expected under several current predictive frameworks. First, this association is consistent with the intuitively plausible idea that resources related to the size of the domicile limit reproduction for inhabitants, such that some end up nonreproductive. In *D. aneurae*, a good candidate for such a limiting resource is the feeding substrate, i.e. the phyllode surface within the domicile. Individuals are thought to feed entirely within their domiciles, perhaps due to highly desiccating conditions outside [32] and have never been documented or observed feeding outside; the phyllode surface within mature domiciles is yellow and necrotic compared to fresh green tissue immediately outside (JDJG, pers. obs.). Thus it seems reasonable to suppose that in small domiciles this surface area may limit reproduction. Alternatively, space within the domicile may be a limiting resource.

Second, however, our data are also consistent with the idea that reproductive skew is the result of a “tug of war” between dominants and subordinates, whereby larger resources (whether limiting to reproduction or not) are more difficult to monopolise by any given individual [19,49]. Third, reproductive skew may reflect the extent of “staying incentives” offered by dominants to entice subordinates not to leave the group, predicting that larger incentives are required where each subordinate contributes proportionally less to group productivity, e.g. where better resources can support a larger group in which subordinate effort is diluted. Finally, skew may reflect “peace incentives” to induce subordinates not to escalate conflict [21,22], which predicts lower skew on better resources where subordinates are stronger and thus more likely to mount a challenge to the dominant.

Thus, a small domicile may (1) provide insufficient resources for all potentially reproductive group members to breed, resulting in intra-domicile competition in which the losers become nonreproductive; (2) provide a small enough arena in which one or a few individuals may be effective in monopolising resources, suppressing the reproduction of others; (3) support a small enough group that subordinates do not require a reproductive incentive to stay – i.e. where a subordinate is capable of making a difference to her kin-groups fitness which on its own outweighs the benefits of breeding independently; or (4) support a small enough group that the threat of escalated conflict does not require a peace incentive and the dominant is able to suppress the reproduction of other members. Teasing apart these alternatives will require detailed experiments.

In large domiciles, despite abundant resources single females tended to fare less well than females in groups. It may be that a single female cannot fully utilize the space available in a large domicile (for example, the median total offspring observed across all singleton domiciles was 6 [±IQR 2-10; range 0-26], compared to the median total offspring potentially attainable in the largest 10% of all domiciles, which was 43 [±IQR 10-58; range 0-87]). In large domiciles, single females may incur deleterious costs of construction and maintenance [50]. Under these circumstances, potential extra females may be beneficial as they may share maintenance and/or defence costs without imposing costs of competition [51].

What is the role of nonreproductive females in *D. aneurae?* One possibility is that they act as nonreproductive helpers to related nestmates, in which case *D. aneurae* could be thought of as a cooperative breeder. Controlling for domicile size, though, developing oocyte volume of breeders went down, not up, when nonreproductive females were present in the domicile (Figure 4a). Whilst we reiterate that this correlative result requires experimental data to confirm, the data suggest that nonreproductive individuals may not have a positive effect upon breeders fecundity, or at least on their ovarian status. They may yet have unmeasured positive effects upon survival of offspring via “assured fitness returns” [52,53], or by “lightening breeders load” [e.g. 54] – for example by helping repair domicile damage to maintain humidity [as suggested in 32] or in a hygienic role such as maintaining middens [29]. Alternatively, nonreproductive females may be waiting to inherit a breeding position within the nest [55,56]. Again, separating these possibilities will require experimental data. One potential explanation that may be discounted, though, is that nonreproductive females may simply be callow or teneral individuals that are not yet reproductively mature – nonreproductive females occurred in domiciles of all ages, containing offspring of all stages from eggs to adults, making this possibility unlikely.

Given the apparent negative effect of nonbreeding females upon breeders fecundity, why would breeding females tolerate nonbreeding nestmates? *D. aneurae* tolerates both foreign conspecifics [29] and inquilines of other species [31]; in this context, perhaps tolerance of nonreproductive nestmates is unsurprising. Nevertheless females are at least capable of evicting males after mating [29], although this behaviour may carry costs. Any conflict of interest over acceptance may be asymmetrical; an individual may gain more by staying (or joining) than the residents lose by accepting her [1,57,58]. Alternatively, as above, nonreproductive females may be tolerated because they confer unmeasured benefits to breeders such as helping with domicile repair or performing hygienic functions.

Why would multiple foundresses make a small domicile, when in small domiciles females may be forced to become nonreproductive or to wait to breed – and, so doing, may even reduce productivity of breeders? One possible scenario would be if small domiciles were typically founded singly and then joined by others later, parasitically, after construction [58,59]. Preliminary lab and field data show that, while females typically cooperate from the moment of domicile initiation [27 and JDJG, pers. obs.], a substantial proportion of established domiciles are also joined by additional females (Gilbert & Simpson, MS in prep). Whether joiners tend to be nonreproductive remains to be established, but one intuitively appealing hypothesis for future research is that vagrant (and presumably unrelated) females may join established domiciles parasitically and compete with foundresses, only becoming reproductive if there are enough resources (i.e. if the domicile is large enough).

Another (nonexclusive) possibility is that cofounding females may not know exactly the eventual size of the domicile before they begin construction. Nothing is yet known about interactions at the moment of domicile formation. It is likely that females have partial but not full control over the eventual size of the domicile they build, because of mechanical constraints imposed by the specific phyllodes they choose to tie together. Thus, a female or group of females may end up in a domicile larger or smaller than optimal [see e.g. 60]. Single females may have less control over domicile location, size and shape than groups of cooperating females – increasing numbers of foundresses are associated with reduced variance in the dimensions of phyllodes used to build domiciles (Gilbert and Simpson, MS in prep).

### Future directions

Experimental data would clearly be necessary to test conclusively how within-group fitness is causally affected by competition [e.g. 12] and should now be a priority for research. In addition, data on genetic relatedness and offspring maternity will be essential to evaluate the costs and benefits of social behaviour in *D. aneurae* in the context of kin selection. Within-group relatedness can demonstrably affect reproductive skew [61,e.g. 62] and may be a critical factor determining whether or not thrips benefit from competing with nestmates over reproduction. One analysis demonstrated that *D. aneurae* has a mixture of high and low relatedness within domiciles [30] consistent with their having later been observed both inbreeding and outbreeding [29] and both cofounding and joining (Gilbert & Simpson, MS in prep); preliminary analyses on our study population reveal that relatedness is mostly very high (LA Rollins, JDJ Gilbert, SJ Simpson, unpublished data). Finally, to understand the dynamics of competition and cooperation before, during and after domicile construction, it will be important to determine how single and multiple foundresses choose and exploit nesting sites, how much control they have over domicile size, and the significance and dynamics of nest-joining behaviour.

## DATA ACCESSIBILITY

All relevant datasets are provided as supplementary material

All R code is available at https://github.com/jdjgilbert/thrips.git

## AUTHOR CONTRIBUTIONS

JDJG conceived the study, designed the work, carried out field and lab data collection, performed the statistical analyses and wrote the manuscript. AW collected key field and lab data. SJS helped to conceive the study and to draft the manuscript. All authors gave final approval for publication.

## ACKNOWLEDGEMENTS

We thank Laurence Mound for invaluable assistance with thrips identification, logistics, help in the field and a wealth of general advice, K. Leggett and G. and V. Dowling for assistance with logistics, and J. Field and L. E. Browning for helpful discussions and comments on the manuscript.

